# SARS-CoV-2 Delta variant induces severe damages in the nasal cavity from the first day post-infection in the Syrian hamster model

**DOI:** 10.1101/2024.08.30.610497

**Authors:** Maxime Fusade-Boyer, Adèle Gambino, Laetitia Merle-Nguyen, Audrey Saint-Albin, Ophélie Ando-Grard, Younes Boujedli, Hélène Huet, Meriadeg Ar Gouilh, Sandra Martin-Latil, Bernard Klonjkowski, Nicolas Meunier, Sophie Le Poder

**Affiliations:** UMR VIROLOGIE, INRAE, École Nationale Vétérinaire d’Alfort, ANSES Laboratoire de Santé Animale, Université Paris-Est, 94700 Maisons-Alfort, France; Unité de Virologie et Immunologie Moléculaires (UR892), INRAE, Université Paris-Saclay, Jouy-en Josas, France; Department of Virology, University Caen Normandie, INSERM Dynamicure UMR 1311, CHU Caen, Caen, France

**Keywords:** SARS-CoV-2 variants, nasal cavity, olfactory epithelium

## Abstract

SARS-CoV-2 replication initiates in the nasal cavity and can spread to the lower respiratory tract. However, the early physiopathological events that occur in the nasal cavity after infection remain poorly understood. In this study, we investigated the initial steps of viral infection from 1 day post-infection (dpi) in Syrian hamsters infected with SARS-CoV-2 D614G, Delta and Omicron (BA.1) variants and compared them with animals sacrificed at 4 dpi. While the level of viral replication in the nasal turbinates of the three groups of hamsters was equivalent at 4dpi, the amount of viral RNA at 1dpi was higher in D614G- and Delta-infected animals than in the Omicron group. No difference in viral RNA levels or inflammatory markers in the nasal turbinates was observed between D614G- and Delta-infected animals, except for a significantly higher level of IFN-λ in the Delta group at 1dpi. Additionally, histological analysis revealed a more rapid diffusion of the Delta virus reaching the posterior zone of the nasal cavity at 1dpi inducing significant damage to the olfactory epithelium. At the same time, the D614G and Omicron infections were essentially restricted to the anterior part of the nasal cavity with less damage observed. Consistently, viral replication was already effective in the lungs of all Delta-infected hamsters at 1 dpi, but only in two of the six D614G animals. Our results highlight the importance of studying viral infection in the nasal cavity very early after infection with a spatial approach to better understand the physiopathology of the different SARS-CoV-2 variants.

## Introduction

Since its emergence in 2019^1^, the intense circulation of the severe acute respiratory syndrome coronavirus virus 2 (SARS-CoV-2) led unsparingly to the appearance of numerous variants^2^. This emergency arose from a higher transmissibility and/or the ability to escape the immune response, and became rapidly dominant in the human population. The first example was the SARS-CoV-2 strain containing the spike mutation D614G which rapidly replaced the original Wuhan strain at the beginning of the pandemic in Europe in 2020, due to its increased transmissibility^3^. So far, the World Health Organization (WHO) labelled five variants as variants of concern (VOCs): Alpha, Beta, Gamma, Delta and Omicron^2^. The most remarkable VOCs regarding biological features are the Delta and Omicron variants. Delta variant, which emerged in India in late 2020, was much more transmissible than the previous strains, but also likely more pathogenic since infection was associated with higher hospitalization risk and ICU admission^4–6^. Delta variant also moderately escaped neutralizing antibodies produced after infection with the original strain^7^. Omicron, which was first detected later in November 2021 in South Africa, is presently predominant all over the World. Omicron is highly transmissible and is also able to evade the humoral immune response due to mutations in the spike protein^8,9^. Its constant genetic evolution led to the emergence of numerous subvariants (BA.1, BA.2, XBB, XBB1.5, etc..)^10,11^. In terms of pathogenicity, Omicron induces less severe infection in humans. This decreased pathogenicity was suspected timely, when field epidemiological data from South Africa suggested a lower hospitalization rate following Omicron emergence^12^. Since, many studies showed its inherent decreased pathogenicity compared to previous variants^13–17^. Understanding the underlying mechanisms that explain the differences in the physiopathology of the different variants remains crucial to adapt the therapeutic and prophylactic countermeasures. To this aim, infections of Syrian hamster appear as a robust model mimicking a variety of symptoms observed during human infection (anosmia, breathing difficulties …)^18,19,20^. Using this model, many studies have compared the physiopathology of the main interesting VOCs but most of them focused their attention later than 2 days post-infection. Here, we hypothesized that the very early events of infection occurring in the nasal cavity; may differ according to the different VOCs. We infected hamsters with the D614G, Delta, and Omicron (BA.1) variants and sacrificed them at 1 day and 4 days post-infection (dpi) for viral and pathological analysis. The initial infection events in the nasal cavity differed depending on the infecting VOCs at 1 dpi. The most striking difference was that the Delta variant caused more severe damage to the olfactory epithelium and exhibited a faster virus diffusion into the posterior zone of the nasal cavity and lungs.

## Materiel and Methods

### Viruses

The SARS-CoV-2 D614G strain BetaCoV/France/IDF/200107/2020 was isolated in the beginning of the pandemic in Europe in March 2020 by Dr. Paccoud from La Pitié-Salpétrière Hospital in France. This strain was kindly provided by the Urgent Response to Biological Threats (CIBU) hosted by Pasteur Institute (Paris, France), headed by Dr. Jean-Claude Manuguerra. The SARS-CoV-2 Delta strain (UCN 46, clade 21A, NCBI n° PP942171) was isolated from nasopharyngeal swabs obtained from patients suffering from respiratory infection, suspected of COVID19 and submitted to molecular diagnosis in Caen Hospital in Normandie during the 2021 summer. The SARS-CoV-2 Omicron BA.1 strain (hCoV-19/Netherlands/NH-RIVM-72291/2021, Omicron variant, lineage B.1.1.529) was obtained through the EVAg platform. Viruses were amplified in VERO-E6 cell culture under 5%CO2 at 37°C for D614G and Delta strains, and at 35°C for the Omicron virus.

### Animal experimentation

Syrian golden hamsters were used to investigate pathogenicity of different variants of SARS-CoV-2 virus. Animals were housed in biosafety level 3 facility at the national veterinary school of Alfort. The protocol was approved by the ANSES/EnvA/UPEC Ethics Committee (CE2A16) and authorized by the French ministry of Research under the number APAFIS#25384-2020041515287655. Three groups of twelve 8-week-old males (Janvier’s breeding Centre, Le Genset, St Isle, France) were each infected by one of the following strains: D614G, Delta, or Omicron (BA.1). Infection was performed by intranasal inoculation of 80µl of Dulbecco’s Modified Eagle Medium (DMEM) containing 5.10^3^ TCID_50_ of SARS-CoV-2 under gas anesthesia with isoflurane. As a control group, five animals were intranasally inoculated with 80µl of DMEM. Animals were weighted daily. At 1 and 4dpi, 6 hamsters from each group were necropsied. The right side of the nasal turbinates (NT) and the right cranial lobe of the lungs were collected and stored at −80°C in beaded tubes (MP Biomedicals) for molecular analysis. The left side of the nasal cavity was harvested for histology.

### Viral replication and innate immunity response analysis

Total RNA TRIzol/chloroform extraction was performed from lungs and nasal turbinates (NT) harvested at 1 and 4 dpi. 1mL of TRIzol (Invitrogen, Carlsbad, CA) was added in each sample tube before mixing them three times at 6000 rpm for 10s in a bead beater (FastPrep MP Biomedicals). Samples were then incubated at room temperature for 5 min before centrifugation for 5min at 12000g at 4°C. Supernatant was collected and 200 µL of chloroform were added before a shaking step for 15s and a room temperature incubation for 15min. Each tube was then centrifuged for 15min at 16 000g and 4°C. About 300-350µl of the aqueous phase of each sample were collected and mixed with 500µL of isopropanol for 15s before incubation at room temperature for 10min. The tubes were then centrifuge at 16 000g for 10min at 4°C. The supernatant was removed and the pellet was washed two times with 75% ethanol before drying at room temperature for 5min and being eluted in 50µl of RNAse free water.

Viral replication and host genes expressions were assessed in the NT and in the lungs by using the QuantitTect SYBR Green RT-PCR kit (Qiagen) performed in 96-well plates in a final volume of 20µl on a LightCycler 96 (Roche, Mannhaim, Germany) according to the manufacturer’s instructions. For each PCR reaction, 100ng of RNA was used and quantification was made using the 2^-Δ*CT*^method and the geometric means of two housekeeping genes (Actin Beta and RPS6KB1). The primers used for genes amplifications are listed in S1 Table.

### Immunohistochemistry analysis

The immunohistochemistry analysis of the olfactory mucosa tissue sections was performed as described previously^21^. Briefly, animal hemi-heads were fixed for 3 days at room temperature in 4% paraformaldehyde (PFA) and decalcified in Osteosoft (Merck Millipore; Saint-Quentin Fallavier; France) for 3 weeks. Blocks were cryoprotected in 30% sucrose. Cryo-sectioning (12 µm) was performed to generate coronal sections of the nasal cavity. Sections were stored at −80 °C until use.

For immunohistochemistry, non-specific staining was blocked by incubation with 2% bovine serum albumin (BSA) and 0.1% Triton. Sections were then incubated overnight with primary antibodies directed against SARS Nucleocapsid protein (1/1000; mouse monoclonal; clone 1C7C7; Sigma-Aldrich) and ionized calcium-binding adapter molecule 1 (Iba1) (1/500; rabbit monoclonal; clone EPR16588; Abcam), Fluorescence staining was performed using donkey anti-mouse-A555; donkey anti-rabbit-A488 (1/800; Molecular Probes A-31570; A32790 respectively).

To assess the impact of the VOCs infection on the olfactory turbinates, we focused our measures on coronal sections of nasal turbinates in an anterior part containing mainly respiratory epithelium and in a posterior part of the nasal cavity, containing the NALT and the end of the Steno’s gland. We selected these areas as our previous results had shown that the D614G VOCs infection starts in this chosen anterior zone and is just appearing in the posterior zone^22^.

To assess the level of infection, we scored the presence of the virus in the respiratory epithelium, olfactory epithelium, Steno’s gland epithelium and cellular debris in the lumen of the nasal cavity on SARS-CoV-2 N protein IHC signal from 0 (absence of infection in the observed zone) to 10 (fully infected zone). To assess epithelium damage, we scored the integrity of the respiratory epithelium (RE) and olfactory epithelium (OE) from 0 (no damage) to 10 (fully damaged zone) based on Hoechst staining according to missing nuclei area, irregularity of the epithelium structure and increased distance between nuclei indicating loosening of the epithelium leading to desquamation^22^. We also assessed the presence of cellular debris in the lumen cavity based on Hoechst staining with a score of zero in absence of debris to 10 when the lumen is filled with cellular debris. All quantifications were performed blindly of the VOCs used for infection. Images for all fluorescent IHC were acquired with a Panoramic Scan 150 (3D Histech) and analyzed with the CaseCenter 2.9 viewer (3D Histech).

### Statistical analysis

Analysis was performed using GraphPad Prism 9.1.2 (GraphPad Software). Comparison between groups was performed using Mann-Whitney test for bodyweight data and for viral and host genes expression data. Immunohistochemistry scores were statistically analyzed using a two-way ANOVA followed by a Tukey’s multiple comparisons test. For all statistical analysis, p-values less < 0.05 were considered significant. Multivariate statistical analysis on RNA levels in nasal turbinates was achieved using Principal Component Analysis (PCoA) with R software. Detailed information on statistical test used, sample size and p-value are provided in the figure captions.

## Results

### 1/ Rapid diffusion of the Delta variant to lungs from day 1 post-infection

Three cohorts of 12 Syrian hamsters were infected intranasally with 5.10^3^ TCID_50_ of either D614G, or Delta or Omicron (BA.1) variants. For each group, six animals were sacrificed at 1 dpi and the six remaining at 4dpi (Figure 1A). The weight of the animals was monitored daily, and we observed a similar weight loss in the hamsters infected with the Delta and D614G variants beginning at 2dpi. Throughout the experiment, hamsters infected with the Omicron variant did not experience weight loss (Figure 1B).

**Figure 1.**
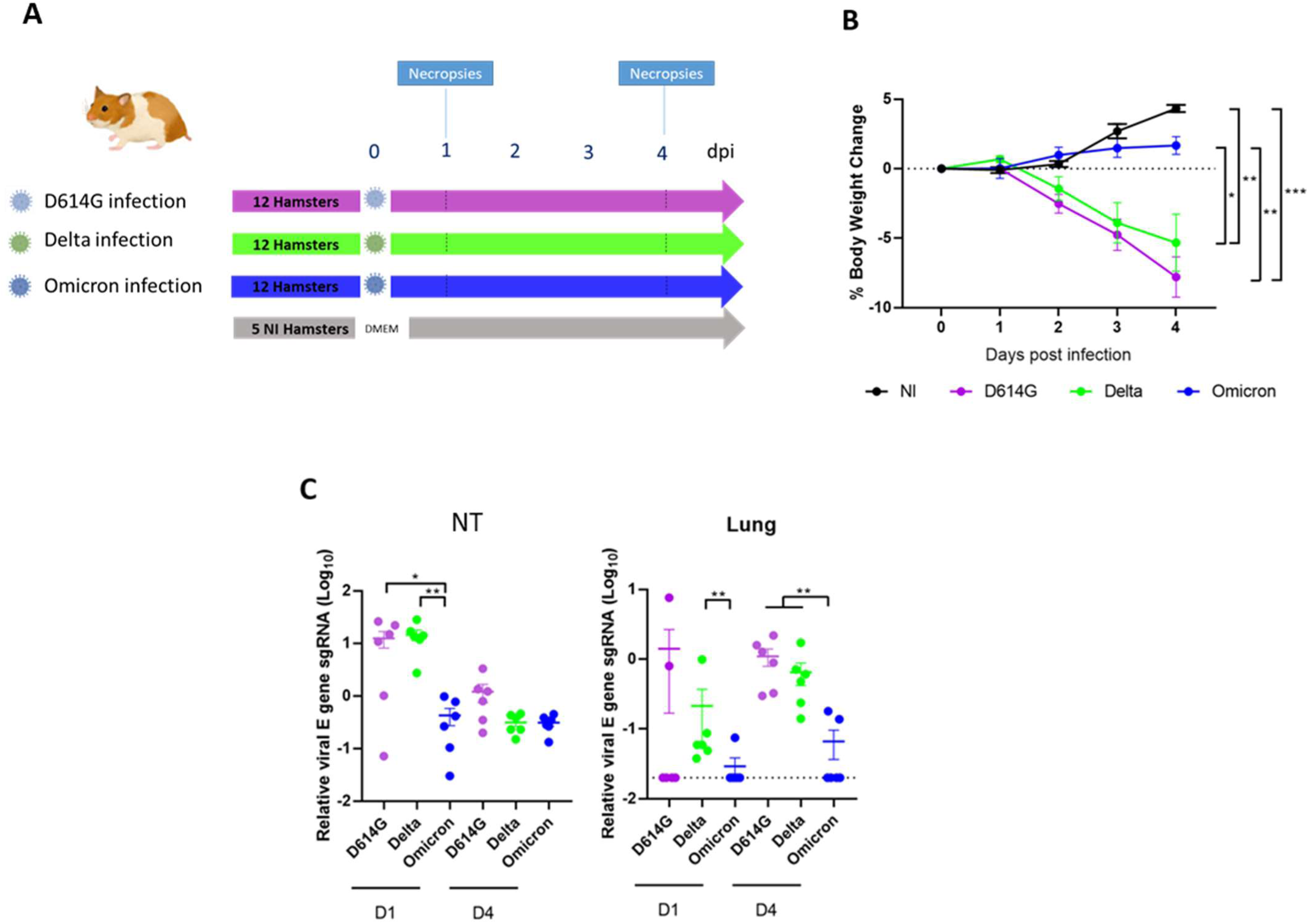
Infection of hamsters with D614G, Delta and Omicron (BA.1) SARS-CoV-2 variants. (**A**) Animal experimental design. Hamsters were infected via intranasal route with 5.10^3^ TCID_50_ of each variant. (**B**) Weight change of animals. Statistical analysis performed with a two-way ANOVA followed by Tukey’s multiple comparison test. (**C**) Viral replication levels in the nasal turbinates (NT) and in the lung at 1dpi (D1) and 4 dpi (D4), determined by RT-qPCR targeting viral subgenomic RNA (sgRNA) relative to housekeeping genes RPS6KB1 and β-actin. Means ± SEM, n=6. Statistical analysis with Mann Whitney tests. *, *P* < 0.05; **, *P* < 0.01; dpi: days post infection; DMEM: Dulbecco’s Modified Eagle Medium, NI: Non-infected

We assessed the level of viral replication in each group of hamsters at 1 and 4 dpi by measuring viral subgenomic RNA targeting the E gene in the nasal turbinates (NT) and in the lungs by RT-qPCR. At 1 dpi, in the NT, levels of SARS-CoV-2 replication were similar with Delta or D614G variants, but about 10-fold lower in hamsters infected with the Omicron variant (Figure 1C) (p<0.05 in comparison with D614G and p<0.001 in comparison with Delta variant). In the lungs, at 1dpi, all the hamsters infected with Delta variant displayed an effective viral replication, whereas this was observed in only 2 of the 6 hamsters in the D614G variant infected group and in only 1 of the 6 hamsters in the Omicron variant infected group.

At 4 dpi, there was no statistically significant difference in viral levels among the three infected groups in the NT. The lungs of hamsters infected with D614G or Delta variants showed similar levels of replication, while in those infected with Omicron variant viral replication was detected in only 2 of 6 hamsters and was 10-fold lower.

### 2/ High elevation of inflammatory markers in the nasal turbinates of D614G and Delta infected animals from day 1 post-infection

Next, we assessed the innate immune response and inflammation triggered by the three SARS-CoV-2 variants by measuring mRNA expression of different targets by RT-qPCR at 1 and 4 dpi in the nasal turbinates (Figure 2).

**Figure 2.**
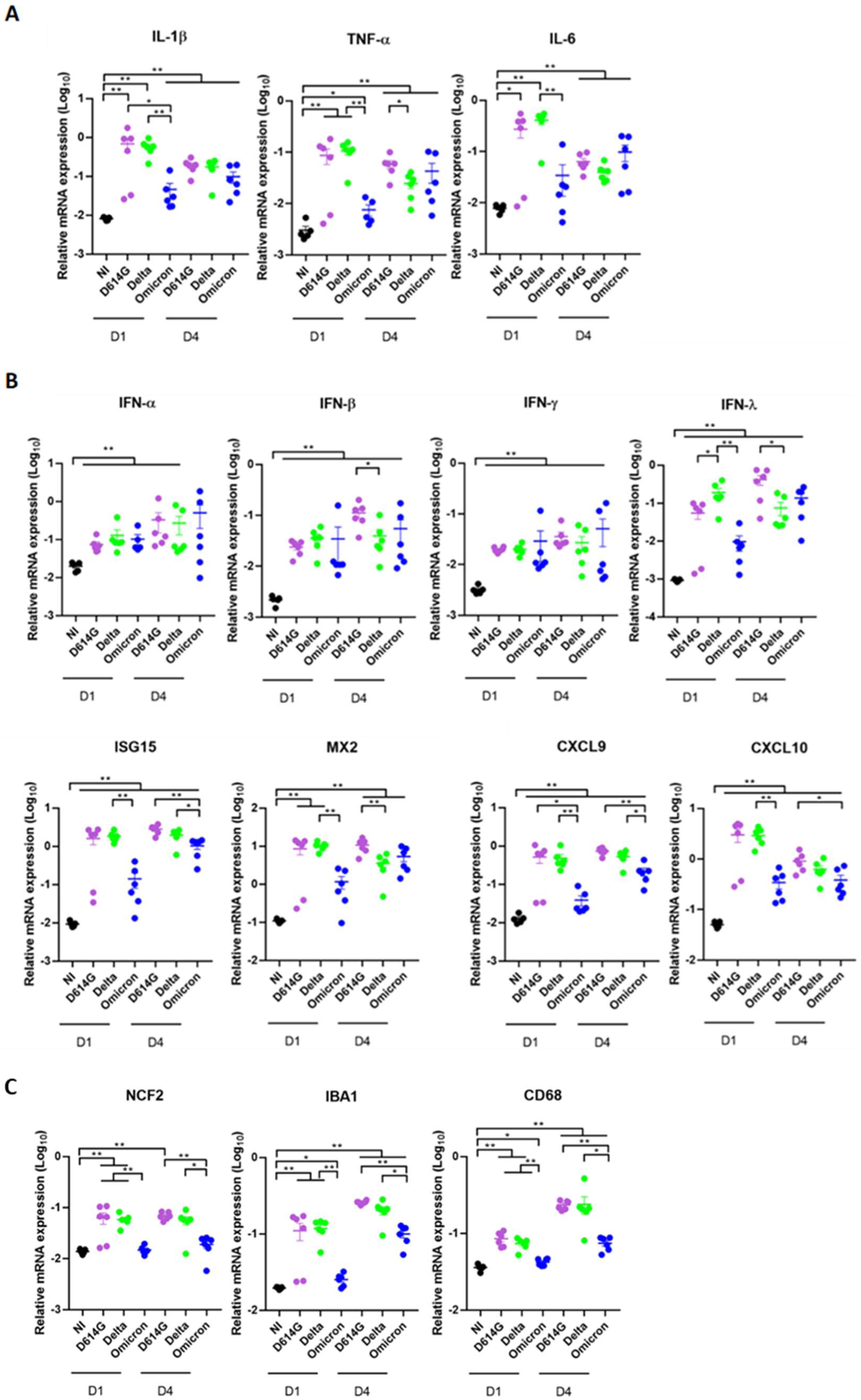
Increase of inflammation markers in the olfactory turbinates of infected hamsters starts as early as 1dpi. mRNA relative expression at 1dpi (D1) and 4dpi (D4) in the olfactory turbinates from hamsters infected with D614G, Delta and Omicron (BA.1) variants **(A)** of inflammatory cytokines (IL-1β, TNF-α, IL-6). **(B)** of markers from the IFN pathway (IFN-α, IFN-β, IFN-γ, IFN-λ, ISG15 MX2) and leucocytes chemo attractive cytokines (CXCL9, CXCL10). **(C)** of neutrophils (Ncf2) and resident / circulating macrophages (Iba1, CD68). Results are expressed as means ± SEM, n=6, Statistical analysis were performed using Mann Whitney tests. *, *P* < 0.05; **, *P* < 0.01. NI=Non-infected

We first studied mRNA expression of the main cytokines involved in inflammatory pathways (Il-1β, TNF-α, IL-6) (Figure 2A). The highest mRNA expression levels of the three main pro-inflammatory cytokines (Il-1β, TNF-α and IL-6) were similarly observed in hamsters infected with D614G and Delta VOC at 1dpi. Hamsters infected with Omicron displayed a 10-fold lower mRNA expression level of these cytokines at 1dpi. At 4 dpi, the expression pattern was similar in the three groups of infected animals, except a lower expression of TNF-α in the Delta group. Then, we measured the mRNA expression of different interferons and ISGs in the NT of infected and non-infected animals at 1 and 4dpi (Figure 2B). The increase expression of IFN-α, IFN-β and IFN-γ was equivalent in all the infected groups at 1 and 4dpi compared to non-infected animals. However, IFN-λ, ISG15 and Mx2 expression levels in infected animals were about 1000-fold higher than in controls and significant differences were observed, especially between Delta and Omicron variants at 1dpi. Interestingly, IFN-λ mRNA expression was different between D614G and Delta infected animals, with a higher level in Delta animals (p<0.05). For these three genes, their expression was about 10-fold lower in hamsters infected with Omicron with a p-value< 0.01 between Omicron and Delta. At 4 dpi, the levels of IFN-λ, ISG15 and Mx2 were still higher than in the non-infected animals. The differences in mRNA expression between the three infected groups were less marked, as these levels tended to increase in the Omicron group compared to 1dpi, whereas they decreased in the Delta group, particularly IFN-λ and Mx2.

Next, we investigated the mRNA expression of chemoattractive cytokines (CXCL9 and CXCL10) and innate immune cells that we had previously characterized during SARS-CoV-2 infection in the NT^22^. We explored indirectly by RT-qPCR the presence of neutrophils (Ncf2), circulating and resident macrophages (CD68; Iba1 respectively) (Figures 2B and 2C). From 1dpi, the transcript levels of all these markers were significantly upregulated in hamsters infected with the D614G and Delta variants. In Omicron-infected hamsters, no increased mRNA expression was observed for Ncf2 and for the other markers the increase was moderate at 1dpi and 4dpi.

We performed the same analysis in the lungs of all hamsters (Figure S1). At 1 dpi, the difference between infected and control animals was more limited than that observed in the nasal turbinates, except for ISG15 and MX2 whose levels were 100-fold higher in the lungs of D614G and Delta animals compared to controls (Figure S1B). Elevation of these ISGS was only 10-fold higher for Omicron infected animals. At 4dpi more markers were elevated in all infected animals. Again, significant differences (p<0.01) were observed between Omicron and the other variants infected animals for many markers (TNF-α, IL-10, IFN-λ, CXCL9, CXCL10, Ncf2, and CD68).

### 3/ Correlation between viral replication and inflammatory markers at 1dpi in the nasal turbinates

As we had noticed differences in the levels of viral replication and inflammatory transcripts between infected animals within the same group, we sought to establish a potential correlation between these parameters.

At 1dpi in the nasal turbinates, we found strong correlations between the level of viral transcripts and IL-1β, TNF-α, IL-6, IFN-λ, MX2, ISG15, CXCL9, CXCL10, Ncf2, Iba1 and CD68 (Figure 3A). We observed no correlation at 4 dpi, which is consistent with the similar levels of the virus and mRNA inflammatory markers among the animals. Finally, PCoA analysis displayed clear separated group of Omicron hamsters from the two other groups (p=0.003) but not the Delta from the D614G (Figure 3B).

**Figure 3.**
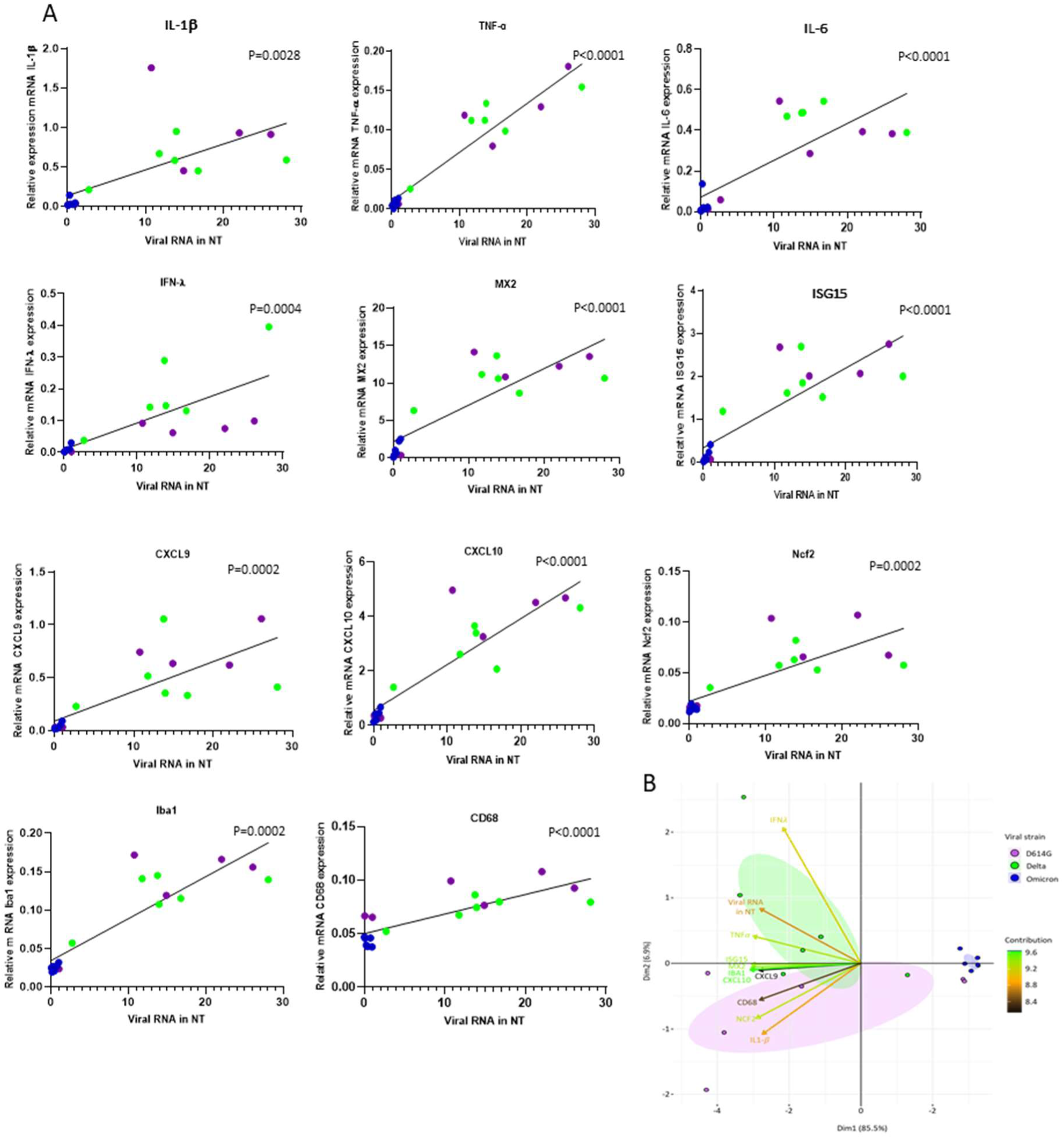
Correlation between viral RNA level and inflammatory markers at 1dpi in the nasal turbinates. (A) Spearman t-test between mRNA relative expression of Il-1β, TNF-α, IL-6, IFN-λ, MX2, ISG15, CXCL9, CXCL10, Ncf2, Iba1, CD68 and the viral level quantified in the nasal turbinates at 1dpi. Significant p-values are indicated **(B)** PCoA biplots of RNA levels in nasal turbinates. Each dot represents a separate animal, colored according to the VOCs. Ellipses represent a confidence interval of 95%. Arrows are colored according to the contribution of the corresponding gene to the first two components of PCoA. The omicron group differs significantly from the others (permutational multivariate analysis of variance with adonis, P < 0.005).

### 4/ The Delta variant causes more damage in the nasal cavity at the early stage of infection

As we observed the main differences between the three groups of hamsters by molecular analysis in the nasal turbinates at 1dpi, we conducted histological examinations of the nasal cavity at this time point. The presence of the virus and the resulting damages were examined in both the anterior and posterior regions of the nasal turbinates (Figure 4A) that we previously observed to be infected at a high and low level respectively at 1 dpi with the D614G variant^22^. In the anterior zone, the level of infection in the respiratory epithelium (RE), which is predominant in this zone, was similar among all three variants (Figures 4A and D). However, the damages to this epithelium were more significant in animals infected with the Delta variant, with more cellular debris in the lumen. The olfactory epithelium (OE) of the anterior zone was equally infected in D614G and Delta animals. However, Omicron animals exhibited significantly lower infection rates in this area (p <0.001 compared to Delta).

**Figure 4:**
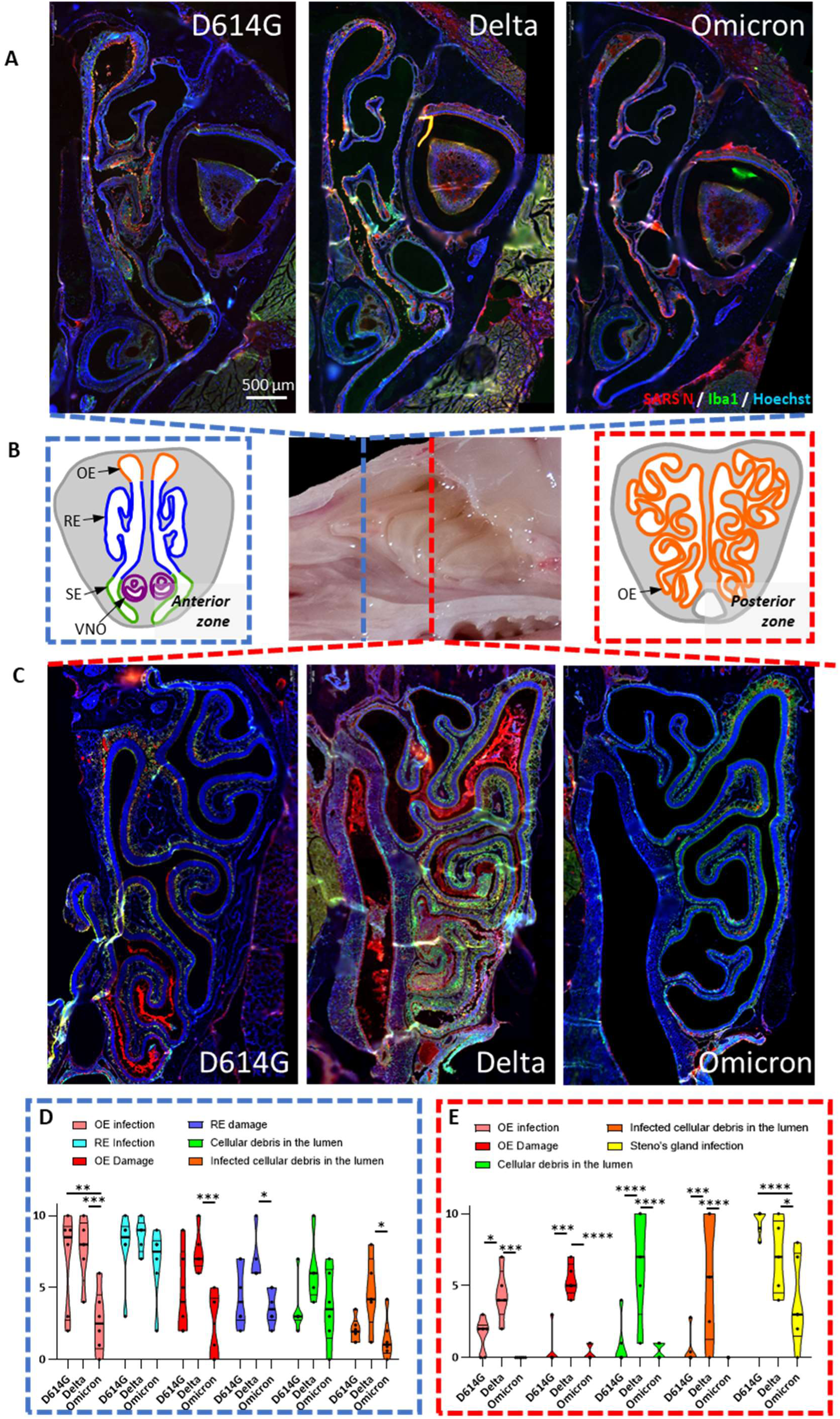
Delta variant induces severe histological damage in the nasal cavity from day 1 post-infection. Hamsters infected with D614G, Delta or Omicron (BA.1) variant were sacrificed at 1 dpi for histological analyses of the nasal cavity and immunostaining of N SARS-CoV-2 antigen (Red) and Iba1 cells (Green). Representative images of the anterior (**A**) or posterior (**C**) zones of the nasal cavity (**B**). Scores of infections and damages induced in anterior zone (**D**) and posterior zone (**E**). Mean ± SEM, n=6, Mann Whitney; *P< 0.05; **P< 0.01, ***P< 0.001, ****P< 0.0001 OE: Olfactory Epithelium, RE: Respiratory Epithelium, VNO: Vomeronasal organ SE: Squamous epithelium

In the posterior zone, histological analysis revealed significant differences with the Delta variant infection. We scored a high presence of the virus in the OE with this variant, in contrast to the D614G variant which exhibited limited infection and Omicron variant no infection at all (Figures 4C and E). Interestingly, the Steno’s gland epithelium was infected by Omicron VOC in this zone. This infection was however lower than with the D614G and Delta VOCs (Fig. 4C). Concomitantly, the olfactory epithelium of animals infected with the Delta variant was greatly altered in this zone with a damage score significantly higher compared to the D614G and Omicron variants (p< 0.001 and p< 0.0001 respectively). We also observed lots of cellular debris filling the lumen of the nasal cavity in these animals (p< 0.0001). In hamsters infected with D614G and Omicron variants, lesions in the nasal cavity were more discreet in this zone, with better integrity of the OE and fewer cellular aggregates in the lumen of the nasal cavity.

## Discussion

Acute SARS-CoV-2 infections lead to different clinical outcomes in humans, from asymptomatic infection to severe respiratory distress and death. Several factors contribute to the diverse range of symptoms, with the infecting SARS-CoV-2 variant being one of them. Omicron and its subvariants cause milder disease than previous variants like D614G and Delta ^13,17^. On the contrary, the number of severe clinical cases was highest during Delta variant infection^5,23^. The Syrian golden hamsters model has been useful to understand the variant pathophysiology differences. Consistent with previous studies^14^, we observed in infected hamsters a low pathogenicity of the BA.1 Omicron variant compared to D614G or Delta variants, with no change of body weight and a moderate tropism to the lower respiratory tract (Figure 1).

For the Delta variant, most studies in hamsters did not reveal any differences compared to D614G or ancestral strains in terms of viral replication level or cytokine inductions^24,25^. In our study, we compared for the first time the diffusion of the different VOCs within the nasal cavity starting at 1dpi. Few damages were observed after Omicron infection. With the D614G variant, the infection resulted in a high level of infected cell debris in the anterior zone of the nasal cavity lumen (Figures 4A and 4D). Interestingly, we observed that the Delta variant spread more rapidly, reaching already at 1dpi the posterior zone of the nasal cavity and the lungs, which are far less affected at this stage with the D614G and Omicron BA.1 infections (Figures 1C and 4C). Moreover, the destruction of the olfactory epithelium was significantly greater after infection with the Delta variant at 1 dpi, particularly in the posterior zone, with statically significant differences (Figure 4E). These particular features observed with the Delta variant could be related to an increased viral fusogenicity which has been linked to the P681R mutation in the spike protein^26,27^. This property allows a more efficient cell diffusion through the formation of syncitia of the Delta variant^26^. With a similar kinetic approach in hamsters, Saito *et al*. study suggested that the Delta virus diffuses more rapidly within the lungs compared to the ancestral strain, reaching earlier the lung periphery even though the total viral load was similar to animals infected with the Wuhan strain.

The pathogenicity of the Delta virus in the nasal cavity seems not to be due to an increase viral replication, as the viral loads in NT were similar between D614G and Delta strains (Figure 1C). Other *in vivo* studies also did not demonstrate higher viral replication with Delta infection^28,29,30^. Nevertheless, we observed heterogeneity among D614G animals with two of them harbouring a lower level of viral replication in the NT at 1dpi. Conversely, in accord with the enhanced spread characteristic of the Delta virus, all animals in this group have already attained a high level of virus load in the nasal turbinates.

As, it was demonstrated that the damages in the olfactory epithelium is induced more by the infiltration of the innate immune cells rather than by the viral replication *per se* ^22^, we analysed by qPCR the presence of neutrophils (Ncf2), resident macrophages (Iba1) and circulating macrophages (CD68). We observed significant lower levels of these cell markers in Omicron -infected animals consistent with the few lesions present in the nasal cavity (Figures 2C and 4D). However, we had no differences between Delta and D614G infected hamsters. We also investigated expression levels of cytokines and interferons responses by qPCR. Again Omicron-infected animals displayed lower expression of them especially at 1dpi, in accordance with previous studies^31^. Only IFN-λ level was significantly higher in Delta animals compared to D614G and no other statistically difference was discernible between both groups (Figure 2B). Contrary to IFN type I, IFN-λ expression is restricted to some cell types^32,33^ among them the epithelial cells from the nasal cavity, which are the SARS-CoV-2 primary target cells. Thus, IFN-λ could be a good marker of SARS-CoV-2 replication in NT. Indeed, the highest level of IFN-λ production in the Delta group could be due to the widespread infection within the entire nasal cavity at 1dpi. Altogether, the molecular results indicated strong correlations between viral replication in NT and induction of inflammatory responses at 1 dpi (Figure 3A). Baker *et al* study observed also a correlation between the viral load in human nasal fluids and several inflammatory cytokines at early stage of infection^34^. Consequently, since the viral loads were lower in the Omicron group, the PCoA analysis enabled clear differentiation of these animals from other infection groups. However, it did not distinguish between the D614G and Delta groups, which had similar quantities of virus (Figure 3B).

The first day post-infection may be too late to distinguish cytokines and interferons responses between D614G and Delta strains. By this time, the infection and the destruction of the OE in the anterior zone is already engaged in D614G animals, triggering the inflammation process (Figure 4A). In Calu-3 cells, infection with the original SARS-CoV-2 strain induces the expression of type I and III IFNs along with the expression of ISGs from 6h post-infection and reaches significant levels from 12h onwards^35^. Then, it would be interesting to assess the induction of the innate immune response *in vivo* at an earlier stage, prior to 24 hours post-infection. This could enable to better investigate the timing of inflammation stimulation and the differences between Delta and D614G variants.

Overall, our results indicate that studying early stages of the infection is very informative to decipher the physiopathology of SARS-CoV-2 variants, and to distinguish their virulence especially when their differences are modest. The SARS-CoV-2 is evolving with the regular appearance of new Omicron subvariants. Some of them (BA.5, BA.2.75 and BA.2.86)seem to be regaining virulence in comparison to BA.1^36,37,38,39^ and it remains necessary to continue to assess the potential pathogenicity of newly emerging variants in order to possibly adapt the public health countermeasures.

## Acknowledgment

We would like to thank all the people from the PRBM platform of ENVA who helped us in the BSL3 animal facility and Alice Peron for technical help.

## Authors contribution

Conceptualization SLP, MFB, NM and AG Investigation: MFB, AG, LMN, ASA, OAG, YB, HH, MAG, SML, BK, SLP, NM Formal analysis: SLP, MFB, NM and AG Writing: SLP, MFB and AG with input from all authors.

## Avaibility of data and material

The data that support the findings of this study are available from the corresponding author, SLP, upon reasonable request. Not applicable for material. Raw data have been deposited at the 4TU.ResearchData databank (https://data.4tu.nl) and are available under the accession number <u>doi.org/10.4121/a4563fc2-5feb-48eb-b195-3ac4ccc99fe9</u>.

---

**S1 Table.**
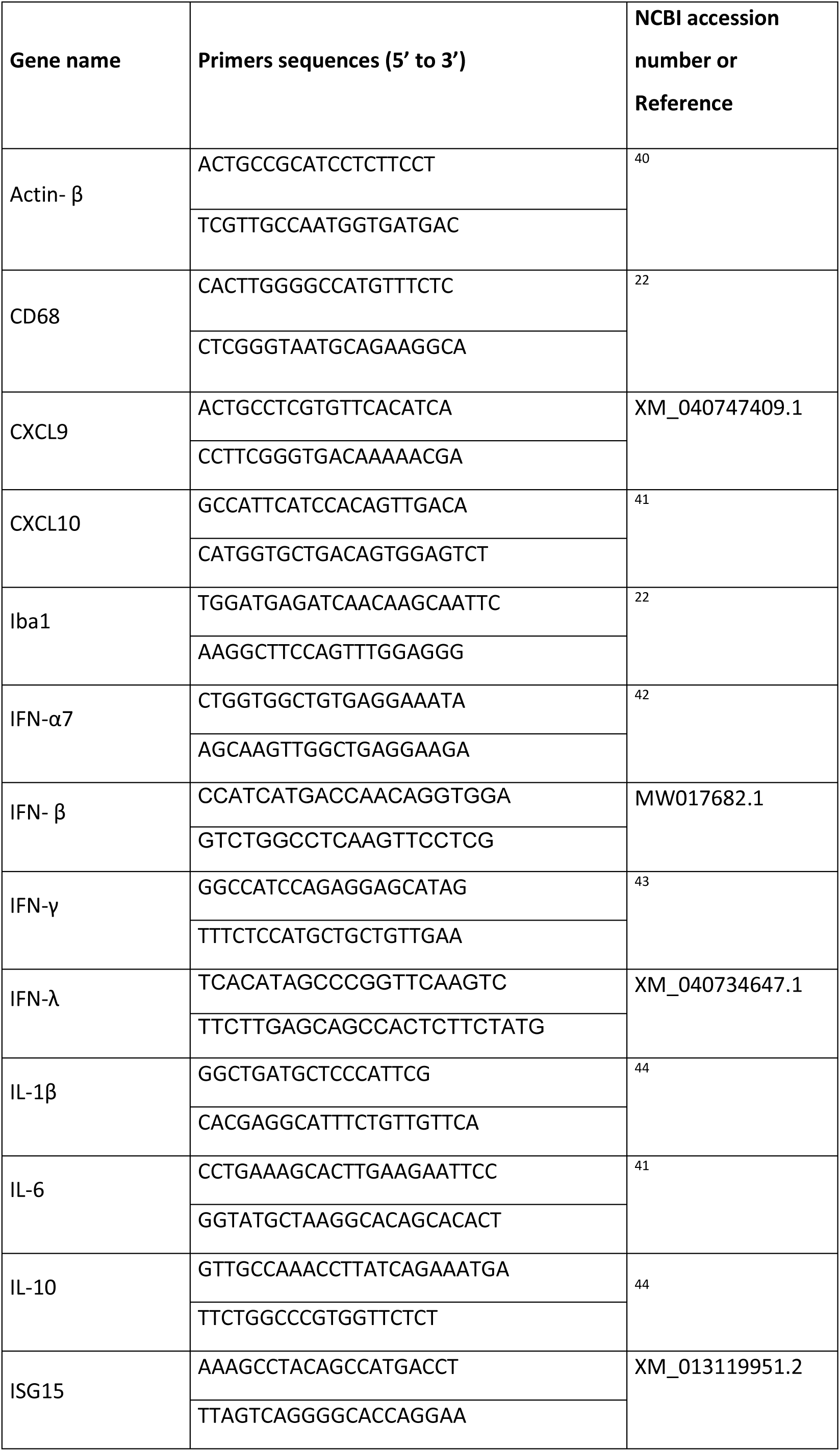
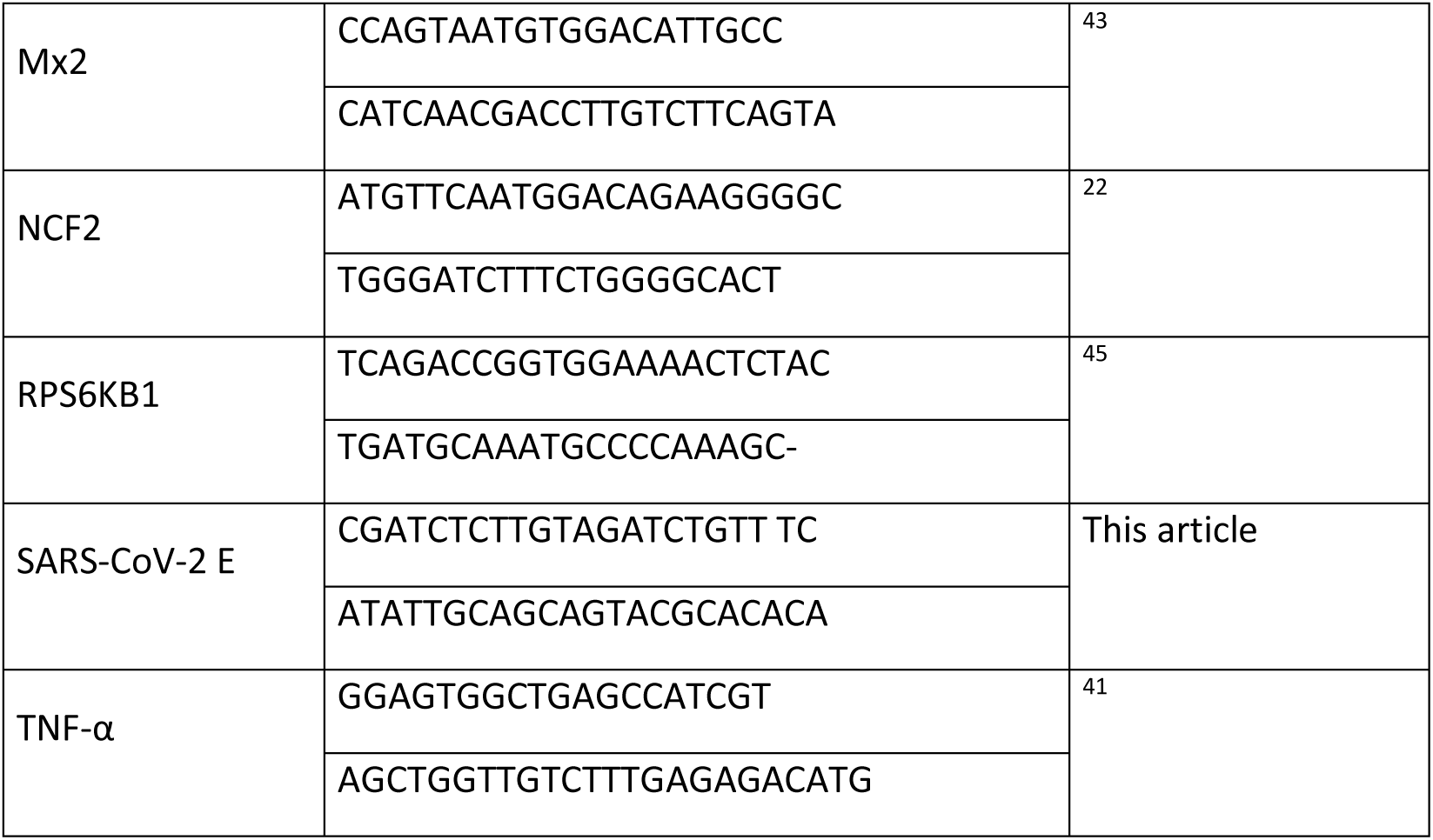
List of primers.

**Figure S1.**
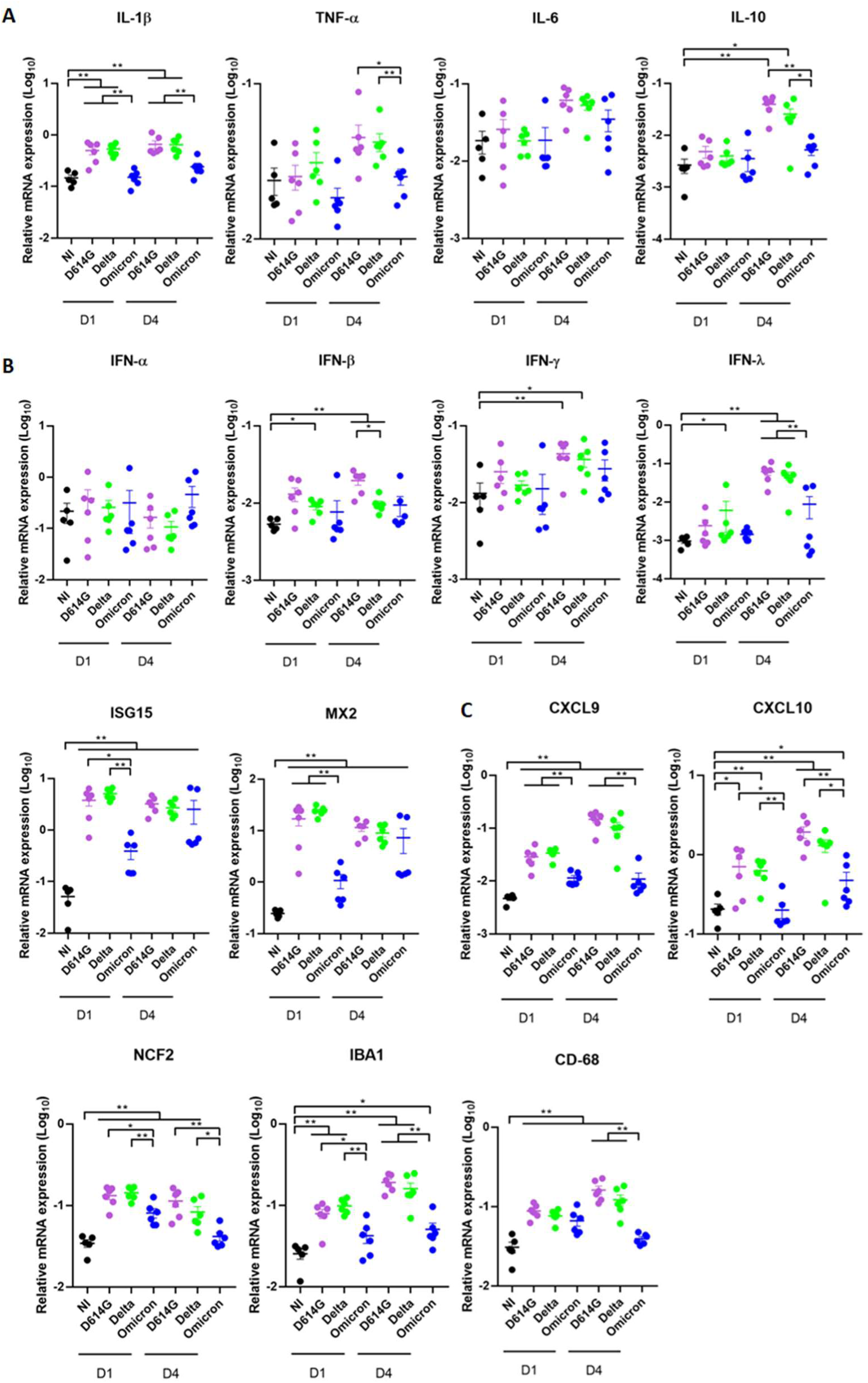
Inflammation markers in the lungs of infected hamsters. mRNA relative expression at 1dpi (D1) and 4dpi (D4) in the lungs from hamsters infected with D614G, Delta and Omicron (BA.1) variants **(A)** of inflammatory cytokines (IL-1β, TNF-α, IL-6). **(B)** of markers from the IFN pathway (IFN-α, IFN-β, IFN-γ, IFN-λ, ISG15 MX2). **(C)** of leucocytes chemoattractive cytokines (CXCL9, CXCL10), neutrophils (Ncf2) and macrophages (Iba1, CD68). Results are expressed as means ± SEM, n=, Statistical analysis were performed using Mann Whitney tests. *, *P* < 0.05; **, *P* < 0.01. NI=Non-infected

## References

1. Wu, F., Zhao, S., Yu, B., Chen, Y.-M., Wang, W., Song, Z.-G., Hu, Y., Tao, Z.-W., Tian, J.-H., Pei, Y.-Y., et al. (2020). A new coronavirus associated with human respiratory disease in China. Nature 579, 265–269. 10.1038/s41586-020-2008-3.

2. WHO (2021). https://www.who.int/activities/tracking-SARS-CoV-2-variants.

3. Zhou, B., Thao, T.T.N., Hoffmann, D., Taddeo, A., Ebert, N., Labroussaa, F., Pohlmann, A., King, J., Steiner, S., Kelly, J.N., et al. (2021). SARS-CoV-2 spike D614G change enhances replication and transmission. Nature 592, 122–127. 10.1038/s41586-021-03361-1.

4. Callaway, E. (2021). Delta coronavirus variant: scientists brace for impact. Nature 595, 17–18. 10.1038/d41586-021-01696-3.

5. Twohig, K.A., Nyberg, T., Zaidi, A., Thelwall, S., Sinnathamby, M.A., Aliabadi, S., Seaman, S.R., Harris, R.J., Hope, R., Lopez-Bernal, J., et al. (2022). Hospital admission and emergency care attendance risk for SARS-CoV-2 delta (B.1.617.2) compared with alpha (B.1.1.7) variants of concern: a cohort study. Lancet Infect. Dis. 22, 35–42. 10.1016/S1473-3099(21)00475-8.

6. Lin, L., Liu, Y., Tang, X., and He, D. (2021). The Disease Severity and Clinical Outcomes of the SARS-CoV-2 Variants of Concern. Front. Public Health 9, 775224. 10.3389/fpubh.2021.775224.

7. Mlcochova, P., Kemp, S.A., Dhar, M.S., Papa, G., Meng, B., Ferreira, I.A.T.M., Datir, R., Collier, D.A., Albecka, A., Singh, S., et al. (2021). SARS-CoV-2 B.1.617.2 Delta variant replication and immune evasion. Nature 599, 114–119. 10.1038/s41586-021-03944-y.

8. Willett, B.J., Grove, J., MacLean, O.A., Wilkie, C., De Lorenzo, G., Furnon, W., Cantoni, D., Scott, S., Logan, N., Ashraf, S., et al. (2022). SARS-CoV-2 Omicron is an immune escape variant with an altered cell entry pathway. Nat. Microbiol. 7, 1161–1179. 10.1038/s41564-022-01143-7.

9. Saxena, S.K., Kumar, S., Ansari, S., Paweska, J.T., Maurya, V.K., Tripathi, A.K., and Abdel-Moneim, A.S. (2022). Characterization of the novel SARS-CoV-2 Omicron (B.1.1.529) variant of concern and its global perspective. J. Med. Virol. 94, 1738–1744. 10.1002/jmv.27524.

10. Sarkar, A., Omar, S., Alshareef, A., Fanous, K., Sarker, S., Alroobi, H., Zamir, F., Yousef, M., and Zakaria, D. (2023). The relative prevalence of the Omicron variant within SARS-CoV-2 infected cohorts in different countries: A systematic review. Hum. Vaccines Immunother. 19, 2212568. 10.1080/21645515.2023.2212568.

11. Tamura, T., Ito, J., Uriu, K., Zahradnik, J., Kida, I., Anraku, Y., Nasser, H., Shofa, M., Oda, Y., Lytras, S., et al. (2023). Virological characteristics of the SARS-CoV-2 XBB variant derived from recombination of two Omicron subvariants. Nat. Commun. 14, 2800. 10.1038/s41467-023-38435-3.

12. Ledford, H. (2021). How severe are Omicron infections? Nature 600, 577–578. 10.1038/d41586-021-03794-8.

13. McMahan, K., Giffin, V., Tostanoski, L.H., Chung, B., Siamatu, M., Suthar, M.S., Halfmann, P., Kawaoka, Y., Piedra-Mora, C., Jain, N., et al. (2022). Reduced pathogenicity of the SARS-CoV-2 omicron variant in hamsters. Med N. Y. N 3, 262–268.e4. 10.1016/j.medj.2022.03.004.

14. Suzuki, R., Yamasoba, D., Kimura, I., Wang, L., Kishimoto, M., Ito, J., Morioka, Y., Nao, N., Nasser, H., Uriu, K., et al. (2022). Attenuated fusogenicity and pathogenicity of SARS-CoV-2 Omicron variant. Nature 603, 700–705. 10.1038/s41586-022-04462-1.

15. Rissmann, M., Noack, D., van Riel, D., Schmitz, K.S., de Vries, R.D., van Run, P., Lamers, M.M., Geurts van Kessel, C.H., Koopmans, M.P.G., Fouchier, R.A.M., et al. (2022). Pulmonary lesions following inoculation with the SARS-CoV-2 Omicron BA.1 (B.1.1.529) variant in Syrian golden hamsters. Emerg. Microbes Infect. 11, 1778–1786. 10.1080/22221751.2022.2095932.

16. Nyberg, T., Ferguson, N.M., Nash, S.G., Webster, H.H., Flaxman, S., Andrews, N., Hinsley, W., Bernal, J.L., Kall, M., Bhatt, S., et al. (2022). Comparative analysis of the risks of hospitalisation and death associated with SARS-CoV-2 omicron (B.1.1.529) and delta (B.1.617.2) variants in England: a cohort study. Lancet Lond. Engl. 399, 1303–1312. 10.1016/S0140-6736(22)00462-7.

17. Halfmann, P.J., Iida, S., Iwatsuki-Horimoto, K., Maemura, T., Kiso, M., Scheaffer, S.M., Darling, T.L., Joshi, A., Loeber, S., Singh, G., et al. (2022). SARS-CoV-2 Omicron virus causes attenuated disease in mice and hamsters. Nature 603, 687–692. 10.1038/s41586-022-04441-6.

18. Muñoz-Fontela, C., Dowling, W.E., Funnell, S.G.P., Gsell, P.-S., Riveros-Balta, A.X., Albrecht, R.A., Andersen, H., Baric, R.S., Carroll, M.W., Cavaleri, M., et al. (2020). Animal models for COVID-19. Nature 586, 509–515. 10.1038/s41586-020-2787-6.

19. Bryche, B., St Albin, A., Murri, S., Lacôte, S., Pulido, C., Ar Gouilh, M., Lesellier, S., Servat, A., Wasniewski, M., Picard-Meyer, E., et al. (2020). Massive transient damage of the olfactory epithelium associated with infection of sustentacular cells by SARS-CoV-2 in golden Syrian hamsters. Brain. Behav. Immun. 89, 579–586. 10.1016/j.bbi.2020.06.032.

20. Sia, S.F., Yan, L.-M., Chin, A.W.H., Fung, K., Choy, K.-T., Wong, A.Y.L., Kaewpreedee, P., Perera, R.A.P.M., Poon, L.L.M., Nicholls, J.M., et al. (2020). Pathogenesis and transmission of SARS-CoV-2 in golden hamsters. Nature 583, 834–838. 10.1038/s41586-020-2342-5.

21. Bryche, B., Fretaud, M., Saint-Albin Deliot, A., Galloux, M., Sedano, L., Langevin, C., Descamps, D., Rameix-Welti, M.A., Eleouet, J.F., Le Goffic, R., et al. (2019). Respiratory syncytial virus tropism for olfactory sensory neurons in mice. J. Neurochem., e14936. 10.1111/jnc.14936.

22. Bourgon, C., Albin, A.S., Ando-Grard, O., Da Costa, B., Domain, R., Korkmaz, B., Klonjkowski, B., Le Poder, S., and Meunier, N. (2022). Neutrophils play a major role in the destruction of the olfactory epithelium during SARS-CoV-2 infection in hamsters. Cell. Mol. Life Sci. CMLS 79, 616. 10.1007/s00018-022-04643-1.

23. Lin, L., Liu, Y., Tang, X., and He, D. (2021). The Disease Severity and Clinical Outcomes of the SARS-CoV-2 Variants of Concern. Front. Public Health 9, 775224. 10.3389/fpubh.2021.775224.

24. Liu, X., Mostafavi, H., Ng, W.H., Freitas, J.R., King, N.J.C., Zaid, A., Taylor, A., and Mahalingam, S. (2022). The Delta SARS-CoV-2 Variant of Concern Induces Distinct Pathogenic Patterns of Respiratory Disease in K18-hACE2 Transgenic Mice Compared to the Ancestral Strain from Wuhan. mBio 13, e0068322. 10.1128/mbio.00683-22.

25. Shahbaz, S., Bozorgmehr, N., Lu, J., Osman, M., Sligl, W., Tyrrell, D.L., and Elahi, S. (2023). Analysis of SARS-CoV-2 isolates, namely the Wuhan strain, Delta variant, and Omicron variant, identifies differential immune profiles. Microbiol. Spectr. 11, e0125623. 10.1128/spectrum.01256-23.

26. Saito, A., Irie, T., Suzuki, R., Maemura, T., Nasser, H., Uriu, K., Kosugi, Y., Shirakawa, K., Sadamasu, K., Kimura, I., et al. (2022). Enhanced fusogenicity and pathogenicity of SARS-CoV-2 Delta P681R mutation. Nature 602, 300–306. 10.1038/s41586-021-04266-9.

27. Rajah, M.M., Hubert, M., Bishop, E., Saunders, N., Robinot, R., Grzelak, L., Planas, D., Dufloo, J., Gellenoncourt, S., Bongers, A., et al. (2021). SARS-CoV-2 Alpha, Beta, and Delta variants display enhanced Spike-mediated syncytia formation. EMBO J. 40, e108944. 10.15252/embj.2021108944.

28. Shuai, H., Chan, J.F.-W., Hu, B., Chai, Y., Yuen, T.T.-T., Yin, F., Huang, X., Yoon, C., Hu, J.-C., Liu, H., et al. (2022). Attenuated replication and pathogenicity of SARS-CoV-2 B.1.1.529 Omicron. Nature 603, 693–699. 10.1038/s41586-022-04442-5.

29. Armando, F., Beythien, G., Kaiser, F.K., Allnoch, L., Heydemann, L., Rosiak, M., Becker, S., Gonzalez-Hernandez, M., Lamers, M.M., Haagmans, B.L., et al. (2022). SARS-CoV-2 Omicron variant causes mild pathology in the upper and lower respiratory tract of hamsters. Nat. Commun. 13, 3519. 10.1038/s41467-022-31200-y.

30. Halfmann, P.J., Kuroda, M., Armbrust, T., Accola, M., Valdez, R., Kowalski-Dobson, T., Rehrauer, W., Gordon, A., and Kawaoka, Y. (2022). Long-term, infection-acquired immunity against the SARS-CoV-2 Delta variant in a hamster model. Cell Rep. 38, 110394. 10.1016/j.celrep.2022.110394.

31. Yuan, S., Ye, Z.-W., Liang, R., Tang, K., Zhang, A.J., Lu, G., Ong, C.P., Man Poon, V.K., Chan, C.C.-S., Mok, B.W.-Y., et al. (2022). Pathogenicity, transmissibility, and fitness of SARS-CoV-2 Omicron in Syrian hamsters. Science 377, 428–433. 10.1126/science.abn8939.

32. Lozhkov, A.A., Klotchenko, S.A., Ramsay, E.S., Moshkoff, H.D., Moshkoff, D.A., Vasin, A.V., and Salvato, M.S. (2020). The Key Roles of Interferon Lambda in Human Molecular Defense against Respiratory Viral Infections. Pathog. Basel Switz. 9, 989. 10.3390/pathogens9120989.

33. Ye, L., Schnepf, D., and Staeheli, P. (2019). Interferon-λ orchestrates innate and adaptive mucosal immune responses. Nat. Rev. Immunol. 19, 614–625. 10.1038/s41577-019-0182-z.

34. Baker, J.R., Mahdi, M., Nicolau, D.V., Ramakrishnan, S., Barnes, P.J., Simpson, J.L., Cass, S.P., Russell, R.E.K., Donnelly, L.E., and Bafadhel, M. (2022). Early Th2 inflammation in the upper respiratory mucosa as a predictor of severe COVID-19 and modulation by early treatment with inhaled corticosteroids: a mechanistic analysis. Lancet Respir. Med. 10, 545–556. 10.1016/S2213-2600(22)00002-9.

35. Banerjee, A., El-Sayes, N., Budylowski, P., Jacob, R.A., Richard, D., Maan, H., Aguiar, J.A., Demian, W.L., Baid, K., D’Agostino, M.R., et al. (2021). Experimental and natural evidence of SARS-CoV-2-infection-induced activation of type I interferon responses. iScience 24, 102477. 10.1016/j.isci.2021.102477.

36. Meehan, G.R., Herder, V., Allan, J., Huang, X., Kerr, K., Mendonca, D.C., Ilia, G., Wright, D.W., Nomikou, K., Gu, Q., et al. (2023). Phenotyping the virulence of SARS-CoV-2 variants in hamsters by digital pathology and machine learning. PLoS Pathog. 19, e1011589. 10.1371/journal.ppat.1011589.

37. Uraki, R., Iida, S., Halfmann, P.J., Yamayoshi, S., Hirata, Y., Iwatsuki-Horimoto, K., Kiso, M., Ito, M., Furusawa, Y., Ueki, H., et al. (2023). Characterization of SARS-CoV-2 Omicron BA.2.75 clinical isolates. Nat. Commun. 14, 1620. 10.1038/s41467-023-37059-x.

38. Tamura, T., Yamasoba, D., Oda, Y., Ito, J., Kamasaki, T., Nao, N., Hashimoto, R., Fujioka, Y., Suzuki, R., Wang, L., et al. (2023). Comparative pathogenicity of SARS-CoV-2 Omicron subvariants including BA.1, BA.2, and BA.5. Commun. Biol. 6, 772. 10.1038/s42003-023-05081-w.

39. Qu, P., Xu, K., Faraone, J.N., Goodarzi, N., Zheng, Y.-M., Carlin, C., Bednash, J.S., Horowitz, J.C., Mallampalli, R.K., Saif, L.J., et al. (2024). Immune evasion, infectivity, and fusogenicity of SARS-CoV-2 BA.2.86 and FLip variants. Cell, S0092-8674(23)01400-9. 10.1016/j.cell.2023.12.026.

40. Thébault, S., Lejal, N., Dogliani, A., Donchet, A., Urvoas, A., Valerio-Lepiniec, M., Lavie, M., Baronti, C., Touret, F., Costa, B.D., et al. (2022). Biosynthetic proteins targeting the SARS-CoV-2 spike as anti-virals. PLOS Pathog. 18, e1010799. 10.1371/journal.ppat.1010799.

41. Marzi, A., Banadyga, L., Haddock, E., Thomas, T., Shen, K., Horne, E.J., Scott, D.P., Feldmann, H., and Ebihara, H. (2016). A hamster model for Marburg virus infection accurately recapitulates Marburg hemorrhagic fever. Sci. Rep. 6, 39214. 10.1038/srep39214.

42. Schountz, T., Campbell, C., Wagner, K., Rovnak, J., Martellaro, C., DeBuysscher, B.L., Feldmann, H., and Prescott, J. (2019). Differential Innate Immune Responses Elicited by Nipah Virus and Cedar Virus Correlate with Disparate In Vivo Pathogenesis in Hamsters. Viruses 11. 10.3390/v11030291.

43. Zivcec, M., Safronetz, D., Haddock, E., Feldmann, H., and Ebihara, H. (2011). Validation of assays to monitor immune responses in the Syrian golden hamster (Mesocricetus auratus). J. Immunol. Methods 368, 24–35. 10.1016/j.jim.2011.02.004.

44. Atkins, C., Miao, J., Kalveram, B., Juelich, T., Smith, J.K., Perez, D., Zhang, L., Westover, J.L.B., Van Wettere, A.J., Gowen, B.B., et al. (2018). Natural History and Pathogenesis of Wild-Type Marburg Virus Infection in STAT2 Knockout Hamsters. J. Infect. Dis. 218, S438–S447. 10.1093/infdis/jiy457.

45. Toth, K., Lee, S.R., Ying, B., Spencer, J.F., Tollefson, A.E., Sagartz, J.E., Kong, I.-K., Wang, Z., and Wold, W.S.M. (2015). STAT2 Knockout Syrian Hamsters Support Enhanced Replication and Pathogenicity of Human Adenovirus, Revealing an Important Role of Type I Interferon Response in Viral Control. PLoS Pathog. 11, e1005084. 10.1371/journal.ppat.1005084.

